# Flipping the Script: *Wolbachia* Favoring Males in a Neotropical *Drosophila*

**DOI:** 10.1101/2024.04.26.591302

**Authors:** Marina Magalhães Moreira, Luísa de Paula Bouzada Dias, Yoan Camilo Guzman, Letícia Sena, João Paulo Pereira de Almeida, Karla Yotoko

## Abstract

As skilful strategists, intracellular endosymbionts, particularly *Wolbachia*, have evolved the ability to induce phenotypes that frequently enhance the fitness of female hosts, often at the cost of male fitness, to ensure the transmission to subsequent host generations. Natural selection plays a pivotal role in this interaction, potentially amplifying, diminishing, or eradicating endosymbionts based on their impact on host fitness. This study investigated the relationship between the *Wolbachia* strain *w*Stv Vi and *Drosophila sturtevanti*, the most abundant Neotropical drosophilid. We combined field sampling and controlled crosses of *Wolbachia*-infected and antibiotic-treated individuals to assess the endosymbiont’s host effects. We found that contrary to initial expectations, *Wolbachia* reduced female fecundity while boosting male fertility, yielding a similar number of offspring in both infected and treated pairs. However, infected females produced fewer larvae when crossed with treated males. A key observation was protogyny in treated but not infected individuals, suggesting *Wolbachia’s* influence on host ontogeny, potentially increasing mating among infected siblings and restoring infected females’ fitness. From these results, we concluded that this whole strategy balanced the fitness of infected and non-infected pairs. In fact, repeated sampling, at the same site, revealed fluctuations in Wolbachia prevalence and high, but not perfect, vertical transmission. Our results indicate that the strategies for persistence in a particular host extend far beyond favoring females. They also imply that environmental factors may favor one group over another in varying circumstances, potentially explaining the observed fluctuations in infection and variable prevalence of *Wolbachia* in *D. sturtevanti* populations.

## 1. Introduction

Intracellular endosymbionts play a fascinating and complex role in evolutionary biology. As a strategic evolutionary adaptation, these endosymbionts have evolved the remarkable ability to induce phenotypes that frequently bolster the fitness of female hosts, often at the expense of males (Vautrin & Vavre, 2009; Landmann, 2019). Natural selection acts upon this interaction in various ways, potentially amplifying, reducing, or even extinguishing endosymbiont populations, depending on how the induced phenotypes affect the fitness of the hosts (Bockoven et al., 2020). Interestingly, the influence of an endosymbiont in phenotypic induction is more substantial than introducing a new allele through mutation. While a new allele may require several generations and combinations with alleles from other *loci*, endosymbionts are inherited as a consistent set of information that interact with host, inducing phenotypic changes more directly and immediately (Law, 1991), thus standing out as powerful agents of evolutionary change (White, 2011).

The Alphaproteobacteria *Wolbachia*, belonging to the order Rickettsiales, are the most prevalent intracellular symbionts of the animal kingdom (Kaur et al., 2021), found in approximately 52% of arthropod species (Weinert et al., 2015). Similar to mitochondrial genomes, *Wolbachia* is vertically transmitted from females to offspring. To ensure their persistence in subsequent generations, certain strains of the bacteria induce reproductive phenotypes that enhance the fitness of infected females (Werren et al., 2008). These phenotypes include parthenogenesis (Stouthamer et al., 1990; Weeks & Breeuwer, 2001; Arakaki et al., 2001); feminization of genetic males (Werren & Beukeboom, 1998; Narita et al., 2007; Negri et al., 2008); male killing, wherein male embryos are selectively eliminated (Dyer & Jaenike, 2004; Kageyama & Traut, 2004); and cytoplasmic incompatibility (CI), leading to reduced offspring viability from matings between infected males and uninfected females (Werren, 1997; Beckmann & Hochstrasser, 2017). Alternatively, *Wolbachia* potentially increases the adaptive value of infected hosts by enhancing the fecundity of infected females (Fry et al., 2004; Weeks et al., 2007), protecting against viral infections (Teixeira et al., 2008; Bruner-Montero & Jiggins, 2023), or providing nutrients that are either not synthesized by the hosts or limited in their environment (Newton & Rice, 2020).

*Drosophila sturtevanti* Duda, 1927 (Diptera: Drosophilidae), a species within the *Drosophila saltans* group of the *Sophophora* subgenus, is regarded as one of the most abundant Neotropical drosophilids (Dobzhansky & Pavan, 1950; Gottschalk et al., 2007; Da Mata et al., 2008; Chaves & Tidon, 2008). In 2006, Miller and Riegler (2006) and Mateos et al. (2006) showed that *D. sturtevanti* harbors *Wolbachia* strains related to *w*Stv, a strain known for inducing cytoplasmic incompatibility (CI) when artificially transferred to *D. simulans* Sturtevant, 1919 (Martinez et al., 2015). The *w*Stv strains are phylogenetically distinct from those infecting other *saltans* species and exhibit high sequence identity with *w*Whi, found in *Lutzomyia shannoni* (Dyar, 1929) (Diptera: Psychodidae) and *L. whitmani* Antunes & Coutinho, 1939 (Miller & Riegler, 2006). A new species within the sturtevanti subgroup, *D. lehrmanae*, was recently described (Madi-Ravazzi et al., 2021). This new species, phylogenetically very close to *D. sturtevanti* (Madi-Ravazzi et al., 2021), also hosts a *Wolbachia* strain named *w*Leh, which is very similar to *w*Stv (Roman, 2022).

In their study, Miller and Riegler (2006) observed that some populations of *D. sturtevanti* were infected by strains related to *w*Stv while others were non-infected. This observation was also reported by Roman (2022), who increased the sampling of populations in South America, particularly in Brazil. She demonstrated that those populations with infected individuals presented prevalence between 30-70%.

In 2022, Zorzato et al. demonstrated that *D. sturtevanti* populations are structured in a North-South direction in terms of mitochondrial haplotypes. These populations exhibit polymorphism and are estimated to have differentiated only about 17,000 years ago, suggesting that *D. sturtevanti* populations either had or still maintain some degree of gene flow, also detected by Trava et al. (2021), in a study involving microsatellites *loci*, justifying the sharing of strains related to *w*Stv.

It is well-established that *Wolbachia* can function as an obligatory, facultative, or even pathogenic endosymbiont (Zug & Hammerstein, 2015). Typically, obligatory endosymbionts are permanently established within a species, which is not the case with the *w*Stv strain (Miller & Riegler, 2006; Roman, 2022). On the other hand, if *w*Stv were pathogenic, natural selection would have already eliminated it from *D. sturtevanti* populations. Therefore, the *w*Stv strains may induce reproductive phenotypes in hosts, contributing to the high incidence of *Wolbachia*-infected individuals observed in South and Central American populations. Given these considerations, our understanding of the *Wolbachia*-host dynamic within the *D. sturtevanti*-*w*Stv system remains as an ongoing study area.

This study unveils the dynamic relationship between *D. sturtevanti* and *Wolbachia*. Through controlled crosses within an isofemale lineage, named st8, originating from a monitored sample site in a preserved fragment of the Atlantic Forest since 2015, we explored critical aspects of this symbiotic interaction. Our investigation spans life cycle stages, assessing fitness factors like fecundity, fertility and, viability. Additionally, we estimated the vertical transmission rate of *w*Stv in *D. sturtevanti*, drawing insights from other isofemale lineages sampled in the same area.

## 2. Methods

### 2.1. Sampling and maintenance of fly lineages

In 2015, we started to investigate the *Wolbachia*-*D. sturtevanti* relationship. Our primary objective was to sample specimens of this species from a preserved fragment of the Atlantic Forest and surrounding areas to assess *Wolbachia* infection. To ascertain whether these specimens harbored *Wolbachia*, we employed traps baited with banana or jackfruit (Dobzhansky & Pavan, 1950), following the methods of Medeiros and Klaczko (1999). We placed traps at five distinct locations, keeping them to remain at those sites for a 48-hour period. The sampled specimens were subsequently transported to our laboratory for detailed analysis. In the lab, we employed CO_2_ to anesthetize the flies, with a particular focus on isolating females from the *saltans* group for further study.

Each female was individually placed in tubes containing a banana-barley culture medium (Supplementary Material 1) maintained at controlled temperature of 23°C and a 12-hour dark/light photoperiod, to promote oviposition and facilitate the generation of F1 males. All experimental procedures in this study were conducted under those conditions.

Using the males obtained in F1, we identified the lineages at the species level as *D. sturtevanti* using the key provided by Markow & O’Grady (2005).

Tubes containing *D. sturtevanti* were maintained as isofemale lineages in the laboratory. Our initial samplings, revealing that all collected isofemales were infected with *Wolbachia*, led us to focus on the preserved Atlantic Forest fragment at the Universidade Federal de Viçosa (UFV) campus. This site, within the UFV boundaries, provides an ideal environment for a longitudinal study due to its accessible and controlled conditions. We conducted additional samplings in 2016, 2019, and 2022 to continually monitor the prevalence of *Wolbachia* at our primary sampling site.

### 2.2. Detection and *Wolbachia* strain identification

We conducted DNA extraction from five female specimens of each isofemale lineage of *D. sturtevanti*, utilising the Wizard Genomic DNA Purification kit (Promega A1120, as detailed in Supplementary Material 2). The integrity of the extracted DNA was evaluated through Polymerase Chain Reaction (PCR), employing *Drosophila* COI primers (Mad-Ravazzi et al., 2021). Samples failing to amplify were excluded from further analysis. Subsequently, *Wolbachia* infection was assessed in the remaining samples by performing standard PCR, utilising wsp226F and wsp588R primers, which are specific to the *Wolbachia* Surface Protein gene (*wsp*) (Arthofer et al., 2009). PCR reactions were prepared in 10 μl volumes, each containing 1.5 mM of each primer, 1x reaction buffer (Promega 5x Green GoTaqG2), 0.04 mM dNTPs, and 0.025 U/μl DNA-Polymerase (Promega GoTaqG2). The PCR protocol involved an initial denaturation at 94°C for 2 minutes, followed by 30 cycles of denaturation at 94°C for 30 seconds, annealing at 67°C for 45 seconds, and elongation at 72°C for 30 seconds. The process was concluded with a final extension step at 72°C for 5 minutes.

We employed both positive and negative controls in our PCR reactions. The positive control consisted of a sample of *D. prosaltans* Duda, 1927 infected with *Wolbachia* [Pro101(Guzman, 2021)]. For negative controls, we used DNA samples from Pro 101 treated with Tetracycline 0.03 % (Guzman, 2021). Additionally, we included a blank control for the DNA extraction process (without biological material) and another blank control for the PCR reaction (without a DNA sample).

The *wsp* gene amplification was checked by applying PCR reaction products in a 1.0% agarose gel electrophoresis and the presence of specific bands (365 bp) was considered evidence of infection and the absence or undetectable presence of bands was considered an indication of no or low infection.

To identify the specific strain or strains of *Wolbachia* present in the *D. sturtevanti* lineage under study, we employed Sanger sequencing. This involved using the multilocus sequence typing (MLST) scheme in conjunction with the *Wolbachia* surface protein (*wsp*) gene analysis, following the methodology established by Baldo et al. (2006). The obtained sequencing traces were meticulously analysed using CodonCode Aligner v 9.0.1 (CodonCode), ensuring accurate and reliable identification of the *Wolbachia* strains.

Given that the *wsp* gene is highly variable and has been used as a tool to differentiate between closely related strains (Miller and Riegler, 2006), we sequenced the *wsp* from other lineages collected in the same region (in 2016, 2019 and 2022) to check whether the population is infected by a single Wolbachia strain or if there are multiple infections.

### 2.3. Antibiotic treatment

From the females collected in 2016 at the preserved fragment of Atlantic Forest in the UFV Campus, we established 11 isofemale lineages of *D. sturtevanti*. Among these, three lineages, st8, st9, and st11, were successfully maintained in the laboratory. However, the st11 lineage lost its infection in mid-2018, and the st9 lineage did not adapt well to laboratory conditions. Therefore, we chose to work with the st8 lineage for antibiotic treatment.

We initiated the treatment by allowing females from the st8 lineage to lay eggs in a banana-barley culture medium (Supplementary material 1) prepared with 0.01% antibiotic Rifampicin (0.1 g/l). We maintained them in this culture medium for four generations. We opted for Rifampicin instead of Tetracycline due to the adverse mitochondrial effects associated with Tetracycline (Ballard & Melvin, 2007), and carefully controlled the Rifampicin dosage to prevent its toxic effects (Lin et al., 2015). Following the treatment, we transferred the fifth generation to an antibiotic-free medium for at least four generations. This recovery period was implemented to mitigate potential confounding factors, such as alterations in gut biota induced by antibiotic treatment (De Crespigny et al., 2008; Li et al., 2016). At each generation of treatment and recovery, we applied the previously described *Wolbachia* detection protocol (Item 2.2) to confirm the treatment effectiveness. From this point forward, we will refer to the untreated individuals from the st8 as belonging to the w+ lineage, and those that underwent Rifampicin treatment as the Rif lineage.

### 2.4. Larval density control

To assess various fitness components, we controlled larval density to obtain adults for subsequent crosses to prevent comparisons between offspring from adults that experienced different nutrition levels during the larval phase that potentially bias the results (Ohba, 1961). Therefore, we transferred groups of 50 first-stage larvae into tubes containing 10 ml of banana-barley culture medium (Snook et al., 2000) (Supplementary material 1) and used the adults for the experiments.

### 2.5. Evaluation of fitness components

#### 2.5.1. Fecundity, fertility and cytoplasmic incompatibility (CI)

After the recovery of the antibiotic treatment (Item 2.3), we set up 14 to 16 replicates for each of the four crosses involving infected (w+) and treated (Rif) st8 lineages: ♀w+ x ♂ w+; ♀w+ x ♂ Rif; ♀Rif x ♂ w+; and ♀Rif x ♂ Rif. Each cross comprised a mating pair of one-day-old virgin individuals. We put each pair on a Petri dish with a culture medium consisting of grape juice (Cattel et al., 2016) (30%), agar (0.8%), and a slender layer of yeast (Supplementary material 1). From the second to the fourteenth day, we transferred each pair to a fresh plate, examined the previous plate for eggs, and counted them. We evaluated the fecundity of the four crosses by comparing the daily eggs laid on the plates. Plates without eggs were discarded. Similarly, fertility was assessed by comparing daily larvae on the remained plates.

To test for Cytoplasmic Incompatibility (CI), we calculated the ratio of Larvae/Eggs produced by each pair per day and compared their values considering the four different crosses (♀w+ x ♂w+; ♀w+ x ♂ Rif; ♀Rif x ♂w+; and ♀Rif x ♂ Rif). The CI is confirmed whenever the pair ♀Rif x ♂w+ presents the lower ratio among the four crosses (Bourtzis et al., 2003).

#### 2.5.2. Total offspring and sex ratio

To evaluate the total offspring and the sex ratio of infected (w+) and treated (Rif) st8 lineages, we conducted crosses involving trios of six-days-old individuals: one female and two males were placed in tubes containing 10 ml of banana-barley culture medium (refer to Supplementary material 1 for details). We set up a total of 25 replicates of w+ and 21 replicates of Rif. In each replicate, adult flies were co-housed for 48 hours to facilitate mating. Post this period, males were removed, and females were allowed an additional 24-hour period in the tubes for oviposition. After the removal of females, each tube was monitored daily for the emergence of adult offspring, which were removed, sexed, and counted. All adults were frozen at −20 °C in 100% alcohol for further analysis. The sex ratio was calculated by determining the ratio of Females / Males that emerged from each tube.

#### 2.5.3. Life cycle

We compared the developmental timeline of infected (w+) and treated (Rif) lineages by recording the number of days until we observed the first egg, first larva (from crosses described in section 2.5.1), first pupa, first female, and first male (from crosses described in section 2.5.2) emerging from the crosses involving only infected (w+) or treated (Rif) st8 lineages.

#### 2.5.4. Longevity

We determined the lifespan of males and females newly emerged from the st8 infected (w+) and treated (Rif) st8 lineages. For that, we placed 10 newly emerged individuals (five males and five females) in tubes containing banana-barley culture medium (Supplementary material 1). We made a total of seven tubes for the infected and 10 for the treated lineage.

We monitored these tubes daily to detect dead individuals, determine their sex, count them, and remove them from the tubes. We transferred the surviving adults to new tubes every five days to prevent generation overlaps. To assess longevity differences between infected (w+) and treated (Rif) st8 lineages, we conducted Log-rank tests (Bland & Altman, 2004) on the entire set of individuals, as well on males and females separately. Significant differences in longevity were depicted using survival courves generated through the Kaplan-Meier method (Kaplan & Meier, 1958). Both Log-rank tests and the survival curves were conducted using the *Survival* package (Therneau, 2015) in R version 4.3.1 (RCoreTeam, 2022).

#### 2.5.5. Vertical transmission rate

Estimating the vertical transmission rate of *Wolbachia* in a given host requires fresh field samplings (Charlat et al., 2004). Therefore, we conduced, in 2019, a new sampling in the Atlantic Forest fragment of the UFV campus. After confirming the species identification in the F1 generation (Section 2.1), we assessed the presence of *Wolbachia* in the newly established *D. sturtevanti* lineages (Section 2.2). To determine the vertical transmission rate, we examined five males and five females from both the F1 and F5 generations of the infected lineages. We conducted these tests using the PCR method described in Section 2.2, with a single-fly protocol for DNA extraction (Supplementary Material 2).

#### 2.5.6. Data analysis

All statistical analyses in this study were conducted using R version 4.3.1 (RCoreTeam, 2022), along with its associated packages. All results, except for those related to longevity data (item 2.5.4), were tested for normality and homoscedasticity. Data exhibiting normality and homoscedasticity were evaluated using ANOVA and are presented with plots depicting each sample mean and standard error when statistically significant. Non-normal data were analysed using the Kruskal-Wallis test (Hollander & Wolfe, 1973). In instances where the Kruskal-Wallis test indicated significance, we further applied the *post-hoc* Dunn test (Dunn, 1964), which compares data from each pairwise cross and applies the Bonferroni correction (*dunn.test*, version 1.3.5; Dinno & Dinno, 2017). Non-normal data were represented using box plots.

## 3. Results

### 3.1. Fly sampling, infection prevalence and *Wolbachia* identification

We have detailed the specifics of all our samplings conducted between 2015 and 2022, including those from the Atlantic Forest fragment at UFV (sample site UFV-1), in Table 1. From these samplings, we established isofemale lineages in the laboratory, initially utilised for identifying species-level and for detecting *Wolbachia* infection. In all samples collected during 2015 and 2016, we observed a 100% prevalence of infection in females. We sequenced the *wsp* gene (Baldo et al., 2006) from the st8 lineage, revealing a sequence identical to the wStv MI strain (accession numbers AY620215.1 and DQ412110.1), previously identified in other *D. sturtevanti* specimens (Mateos et al., 2006; Miller & Riegler, 2006). Additionally, we sequenced the MLST from st8 (GENBANK MW854824 to MW854829). Due to the absence of other alleles (MLST) for comparison with the *w*Stv strain found in our study and those previously reported, we have designated this strain as *w*Stv Vi, reflecting its identification in Viçosa, MG, Brazil. In later samplings, conducted in 2019 and 2022, we tested all isofemale lineages for Wolbachia infection using the methods outlined in Section 2.2 and found that not all were infected (Table 1). Sequencing the *wsp* of a portion of the infected lineages (from samplings 9, 10 and 11, Table 1) revealed that all had the same *wsp* sequence (GENBANK PP540051 to PP540058), suggesting the population under study is infected by a single *Wolbachia* strain.

**TABLE 1:** *D. sturtevanti* samplings in Viçosa (V) at Universidade Federal de Viçosa Campus (UFV), Araponga (A), and Juiz de Fora (JF) in 2015, 2016, 2019, and 2022, showing the number of isofemale lineages (N) and the number of infected lineages (w+).

| Sampling | Date | Sample Site | GPS coordinate | Bait | N | w+ |
| --- | --- | --- | --- | --- | --- | --- |
| 1 | Jan/15 | UFV-1 (V) | 20°45'48"S; 42°51'53"W | Banana | 5 | 5 |
| 2 | Feb/15 | UFV-2 (V) | 20°48'10"S; 42°51'48"W | Banana | 2 | 2 |
| 3 | Apr/15 | Brigadeiro (A) | 20°40'23"S; 42°29'28"W | Banana | 2 | 2 |
| 4 |  | UFV-3 (V) | 20°45'34"S; 42°52'14"W | Banana | 1 | 1 |
| 5 |  | Juiz de Fora (JF) | 21°48'06"S; 43°23'16"W | Jackfruit | 2 | 2 |
| 6 | May/15 | UFV-3 (V) | 20°45'34"S; 42°52'09"W | Jackfruit | 1 | 1 |
| 7 |  | UFV-3 (V) | 20°45'35"S; 42°52'14"W | Banana | 1 | 1 |
| 8 | Aug/15 | UFV-1 (V) | 20°45'48"S; 42°51'53"W | Banana | 10 | 10 |
| 9 | Aug/16 | UFV-1 (V) | 20°45'48"S; 42°51'53"W | Banana | 11 | 11 |
| 10 | Apr/19 | UFV-1 (V) | 20°45'48"S; 42°51'53"W | Banana | 12 | 6 |
| 11 | Apr/22 | UFV-1 (V) | 20°45'48"S; 42°51'53"W | Banana | 12 | 9 |

### 3.2. Antibiotic treatment

The treatment of st8 (item 2.3), conducted with Rifampicin, between February and June of 2018, was successful. From the second generation under antibiotic treatment on, detecting the infection using the protocols described in Section 2.2 was no longer possible. The treated lineage (Rif) remains in the laboratory (January 2024). Periodic checks have confirmed the absence of *wsp* bands in this lineage.

### 3.3. Fitness components

#### 3.3.1. Fertility, fecundity and cytoplasmic incompatibility (CI)

When comparing the four crosses (♀w+ x ♂w+; ♀w+ x ♂Rif+; ♀Rif x ♂w+ and ♀Rif x ♂Rif), we identified significant differences in the daily egg production over the 14-day monitoring period (Kruskal-Wallis χ²=12.974, df=3, p=0.004694). Specifically, the cross involving treated females and infected males (♀Rif x ♂ w+) yielded significantly more eggs than the two crosses involving infected females (♀w+ x ♂ w+ and ♀w+ x ♂ Rif) (Dunn’s test: p= 0.0089; and p = 0.0090, respectively, Fig. 1A).

**FIGURE 1:**
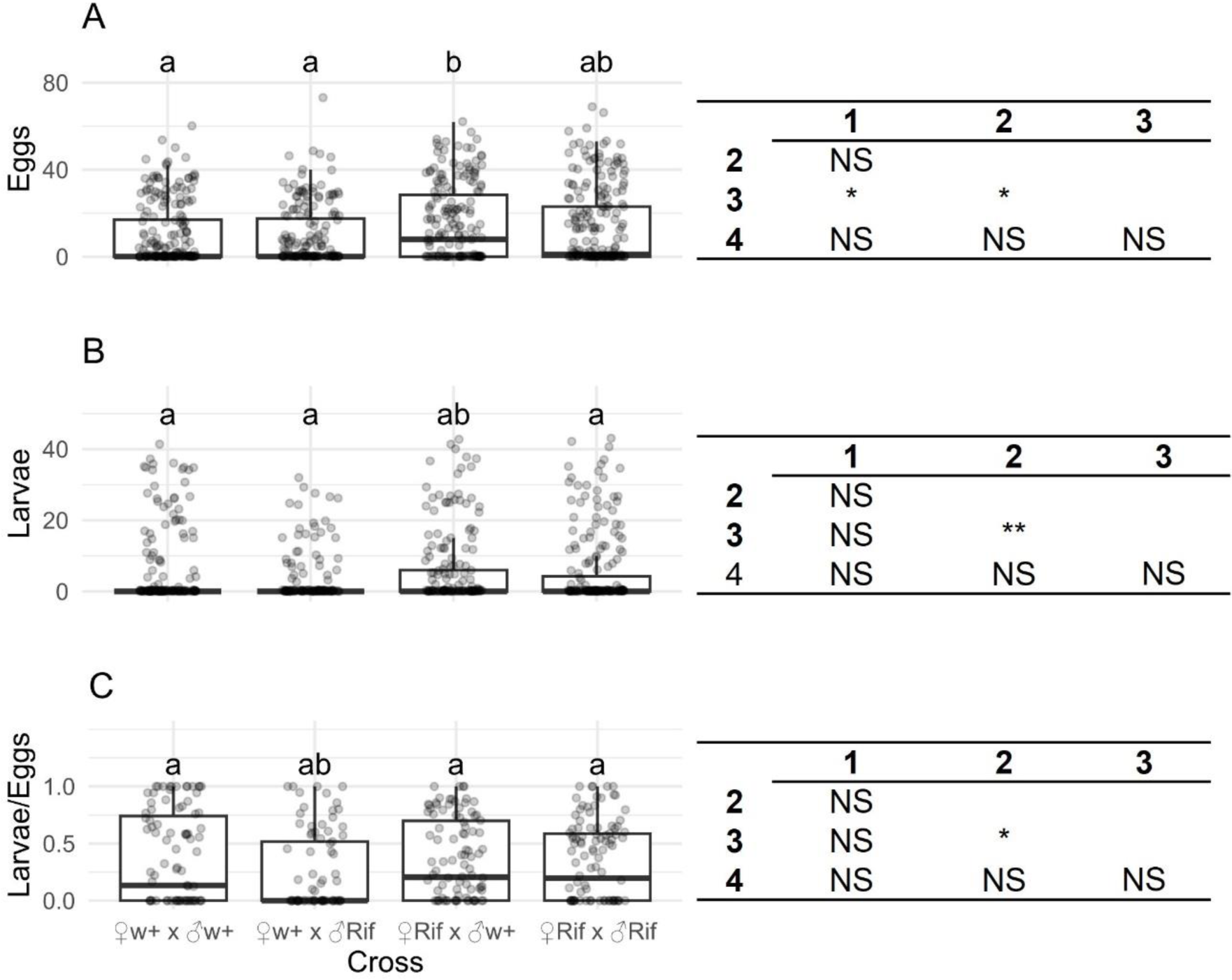
Comparative analysis of reproductive parameters in *D. sturtevanti* crosses involving *Wolbachia*-infected (w+) and antibiotic-treated (Rif) st8 individuals. A. Fecundity, shown as the number of eggs produced per day. B. Fertility, represented by the number of larvae produced per day. C. Egg-to-larvae ratio, indicating the ratio of eggs that resulted in hatched larvae per day. Each dot represents the count of eggs (panel A), larvae (panel B), or the egg-to-larvae ratio (panel C) for each day in each replicate of the respective crosses. Lowercase letters above the bars denote significance groups determined by Dunn’s test. Beside each graph, the significance of the differences between the measures is shown. NS - not significant, *p<0.025, **p< 0.005. The numbers represent the crosses, where: 1 - ♀w+ x ♂w+; 2 - ♀w+ x ♂ Rif; 3 - ♀Rif x ♂w+; 4 - ♀Rif x ♂ Rif.

Regarding larval production, significant differences were also observed among the four crosses (Kruskal-Wallis χ²=14.927, df=3, p=0.00188). In this instance, the ♀Rif x ♂w+ cross yielded significantly more larvae than the ♀w+ x ♂Rif cross (Dunn’s test: p= 0.0006; Fig. 1B).

A significant difference was also observed when comparing the egg-to-larva ratio among the four crosses (Kruskal-Wallis χ²=7.9985, df=3, p=0.04604). Notably, the cross ♀Rif x ♂w+ exhibited a significantly higher egg-to-larva ratio compared to cross ♀w+ x ♂Rif (Dunn’s test: p= 0.0238; Fig. 1C), indicating that the *w*Stv Vi strain does not induce CI in st8.

#### 3.3.2. Total offspring and sex ratio

There was no statistically significant difference in the total number of adults produced by infected (♀w+ x ♂w+) and treated (♀Rif x ♂Rif) lineages (F_1,44_ = 0.4683, p = 0.4973). Furthermore, no statistically significant deviations from the expected 1:1 sex ratio were observed (♀w+ x ♂w+; F_1,48_ = 0.021, p = 0.8855; ♀Rif x ♂Rif; F_1,40_ = 0.2285, p = 0.6352). These results suggest that the *w*Stv Vi strain does not induce selective male-killing in the st8 flies.

#### 3.3.3. Life cycle and longevity

Comparing the crosses (♀w+ x ♂w+ and ♀Rif x ♂Rif), we did not find statistically significant differences in the number of days until the first egg (Kruskal-Wallis χ²=3.6106, df=3, p=0.3067), until the first larva (Kruskal-Wallis χ²=3.4087, df=3, p=0.3328), and until the first adult (Kruskal-Wallis χ²=1.5701, df=1, p=0.2102). However, larvae pupated significantly earlier in the treated lineage (Rif) (Fig. 2).

**FIGURE 2:**
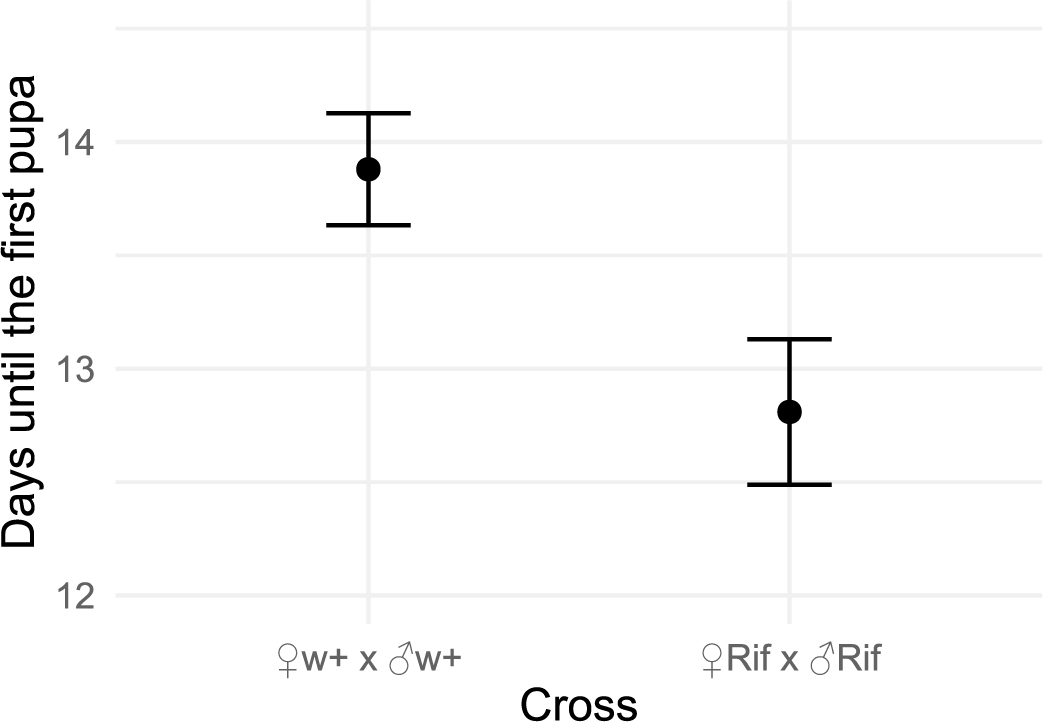
Mean and standard error of the number of days counted from oviposition to the detection of the first pupa in crosses involving infected (w+) and treated (Rif) st8 lineages of *D. sturtevanti* (F1,46 = 7.20, p = 0.01021).

In the crosses involving the infected st8 lineage (♀w+ x ♂ w+), females and males emerged concurrently (Kruskal-Wallis χ²=3.6319, df=1, p=0.05668). Conversely, in the crosses involving the antibiotic-treated st8 lineage (♀Rif x ♂Rif), females emerged significantly earlier than males (Kruskal-Wallis χ²=15.487, df=1, p=8.308e-05) (Fig. 3).

**FIGURE 3:**
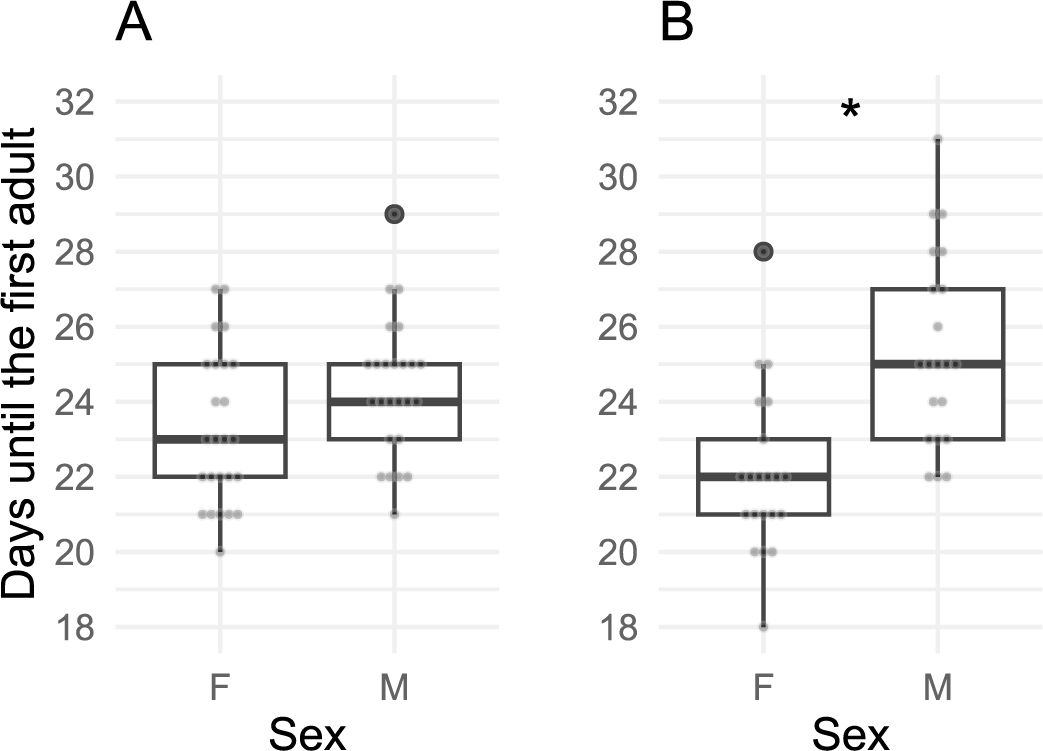
Number of days counted from oviposition to the hatching of the first female (F) and the first male (M) in crosses involving individuals from the st8 lineage of *D. sturtevanti* (A) infected with *Wolbachia* (♀w+ x ♂ w+) and (B) treated with Rifampicin (♀Rif x ♂Rif). The asterisk (*) indicates statistical significance (p < 0.05) inferred by the Kruskal-Wallis test.

We found no significant difference in terms of longevity (days until the death) of w+ and Rif all individuals (p = 0.69), only females (p=0.18) or only males (p=0.39). However, in both lineages, males lived longer than females (p < 0.05), regardless of the infection status (Supplementary material 3).

#### 3.3.4. Vertical transmission rate

In 2019, we sampled 12 *D. sturtevanti* females, of which only six were infected with Wolbachia. The offspring of five out of these six infected isofemales was also infected. The offspring tested was from F1 and F5 which were comprising of 10 individuals in the F1 generation and other 10 in the F5 generation. In the remaining lineage, 18 individuals were infected and two were uninfected: one male at F1 and one female at F5 (90% infected in each generation). These results indicate a high and almost perfect rate of *Wolbachia* transmission within the population at our primary sampling site.

## 4. Discussion

In this study, we aimed to investigate aspects of the *Wolbachia*-host relationship within a lineage of *D. sturtevanti* naturally harbouring *Wolbachia*. We started our research by determining whether *D. sturtevanti* specimens from southeastern Minas Gerais, Brazil, were infected with *Wolbachia* and identifying the infecting strain. Our 2015 sampling confirmed *Wolbachia* infection across all examined lineages (100% prevalence, Table 1). Following this initial approach, we directed our subsequent samplings towards the population at the preserved fragment of Atlantic Forest located within the campus of the Universidade Federal de Viçosa, (Sample Site UFV-1, Table 1).

In 2016, we observed a 100% prevalence of infection within our primary sample site. Among the isofemale lineages established from this sampling, we selected one, named st8, for identifying the *Wolbachia* strain and subsequent crosses. The identified *Wolbachia* strain, sharing the *wsp* (*Wolbachia* surface protein) allele previously noted in Central American lineages (Miller & Riegler, 2006; Mateos et al., 2006) and designated as *w*Stv MI, highlights that *D. sturtevanti* populations are not isolated, suggesting gene flow among geographically disparate populations (Trava et al., 2021; Zorzato et al., 2022). The wsp from other lineages collected at the same site was identical to that found in st8, suggesting the population is infected by a single strain.

Despite earlier reports of naturally uninfected individuals in different populations of *D.sturtevanti* (Miller & Riegler, 2006; Roman, 2022), the widespread *Wolbachia* infection observed until 2016 led us to hypothesise that the *Wolbachia* strain found in st8, termed *w*Stv Vi, might induce reproductive phenotypes like cytoplasmic incompatibility (CI) or confer adaptive benefits to infected females. However, our results failed to support these hypotheses.

Indeed, our experiments involving the st8 lineage yielded an unexpected result: Rifampicin treatment increased egg production (Fig 1A). Such result suggests that the *w*Stv Vi strain may potentially reduce the fecundity of infected st8 females (Fig. 1A), mirroring phenomena observed in more recent symbiotic relationships established through the artificial infection of vectors for the control of infectious diseases (Lau et al. 2022).

Conversely, infected males exhibited higher fertility than treated males, a rare phenomenon previously documented in other insects such as the flour beetle *Tribolium confusum* (Duval, 1868) (Wade & Chang, 1995), the stalk-eyed flies (Diptera: Diopsidae) (Hariri et al. 2002) and pea aphids, *Acyrthosiphon pisum* (Harris) (McLean et al. 2017). Figure 1B indicates that crosses involving a treated female and an infected male (♀Rif x ♂w+) produced significantly more larvae than the other three crosses. This finding is intriguing, because *Wolbachia* infection is known to reduce male fertility (Snook et al. 2000; De Crespigny & Wedell, 2006; Lewis et al., 2010). Also, cytoplasmic incompatibility typically results in the lowest offspring production in crosses involving the pair ♀Rif x ♂w+, where infected males potentially causing reduced egg hatch rates in uninfected females (Biwot et al., 2020).

Furthermore, our results suggest that infected males are more efficient than treated ones at fertilising the eggs of infected females. In Figure 1C, the pair ♀w+ x ♂w+ showed an egg-to-larvae ratio equivalent to the pairs ♀Rif x ♂w+ and ♀Rif x ♂Rif, indicating that mating with w+ males reduced the detriment caused by the reduction of eggs in infected females. Consequently, the pair with the lowest egg-to-larvae ratio consisted of an infected female and a treated male (♀w+ x ♂Rif). Previous work on infected and treated *D. septentriosaltans* Magalhaes & Buck, 1962, another species from the *saltans* group, showed a higher number of inseminations by infected males in both infected and treated females after 48 hours (Guzman, 2021). Similar findings in *D. melanogaster* Meigen 1830 and *D. simulans* (De Crespigny et al., 2006) were interpreted as an adaptive response to mitigate the effects of cytoplasmic incompatibility (CI) in these species. However, neither *D. septentriosaltans* (Guzman, 2021) nor *D. sturtevanti* exhibit CI, necessitating further studies to comprehend the mechanisms that enable greater fertilisation success by infected males. As a result, infected males may exhibit enhanced mating abilities or increased fertilisation skills, aspects that require further exploration in subsequent studies.

Our findings also indicated that larvae from treated pairs (♀Rif x ♂Rif) initiated pupation significantly earlier than those from infected pairs (♀w+ x ♂w+), as depicted in Figure 2. This accelerated development in uninfected flies could confer an advantage, particularly in environments characterized by ephemeral resources and larval crowding, which is typical for *Drosophila* species (Prasad et al., 2001). Moreover, faster pupation leads to quicker maturation into adults (Kumar et al. 2006), potentially allowing these adults earlier access to scarce resources.

The underlying causes of this developmental discrepancy become clearer upon examining the emergence of the first females and males. In treated lineages (Rif), where larvae pupated earlier, females emerged significantly before males (Fig. 3B), providing evidence that earlier pupating larvae led to the emergence of earlier females. This phenomenon, known as protogyny, has been previously observed in species such as *D. ananassae* Doleschall, 1858, *D. malerkotliana* Parshad and Paika, 1965, and *D. melanogaster* (Bharathi et al., 2004; Fellowes et al., 1998). Protogyny is a phenomenon that may have been selected to prevent inbreeding (Seong et al. 2023). In contrast, infected lineages did not exhibit this pattern; instead, infected females emerged simultaneously with their brothers (Fig. 3A). This simultaneity in emergence likely increases the probability of mating between infected siblings, which, according to our results, led to a higher egg-to-larvae ratio compared to the ♀w+ x ♂Rif pair. Such simultaneity could be particularly beneficial in populations where the prevalence of this specific *Wolbachia* strain is low, as it ensures reproductive success for infected females.

Despite the disadvantages on infected female fecundity, the adult production of infected (w+) was equivalent to that of non-infected (Rif) st8 lineages. This led us to believe that the high prevalence observed in 2015/2016 resulted from efficient vertical transmission, which, according to Porter and Sullivan (2023), is a prerequisite for the spread of *Wolbachia*. However, our 2019 sampling revealed a prevalence of only 50% (Table 1). Additionally, of the six isofemale lineages that were infected, only five exhibited perfect vertical transmission. In this context, we also observed that one of the lineages established in the laboratory in 2016, st11, lost its infection without antibiotic treatment, revealing that the 100% prevalence was transient in our sampling site.

The reduction in prevalence observed in 2019 led us to consider the possibility of the infection’s extinction over time. However, our 2022 collection at the exact location revealed a different scenario: only three out of 12 isofemale lineages were uninfected, indicating a resurgence in bacterial prevalence to 75% (Table 1). This fluctuation suggests that bacterial prevalence varies possibly in response to environmental changes that sometimes favor infected flies and, at other times, favor uninfected ones, in ways not measurable in our controlled-laboratory observations. Lindsay et al. (2023) showed that in *D. melanogaster*, *Wolbachia* may offer nutritional advantages in an environment where resources are scarce. Considering that resource availability fluctuates in natural environments, it is plausible to hypothesize that *Wolbachia* prevalence fluctuates in response to that.

Large populations such as those of *D. sturtevanti* (Dobzhansky & Pavan, 1950; Gottschalk et al., 2007; Da Mata et al., 2008; Chaves & Tidon, 2008) suggest significant natural selection acting on slightly varying phenotypes (Wright, 1931). Interestingly, the lineage st8 harbors an endosymbiont that appears to diminish fertility in infected females. However, this disadvantage is counterbalanced by the higher fertility of infected males, as crosses of infected or uninfected pairs results in similar numbers of offspring. If an infected female joins a non-infected group, its endogamous offspring could effectively compete with uninfected females, explaining the absence of protogyny in the offspring of infected pairs, as indicated by our findings. Nevertheless, it is crucial to acknowledge that these interactions only level the fitness of infected and non-infected pairs in our laboratory conditions. This indicates that environmental variables, still to be fully understood, may preferentially favor one group over the other in different habitats, as in other facultative endosymbionts (Correa & Ballard, 2016).

These natural selection fine-tuning could account for the diverse infection rates as well as the non-occurrence of infection in natural *D. sturtevanti* populations, as noted in the studies by Miller and Riegler (2006) and Roman (2022), along with the fluctuations we noted in our primary sample site.

## Acknowledgments

This work is part of Marina Moreira’s Master’s dissertation, supported by a CNPq scholarship and was partially funded by the Coordenação de Aperfeiçoamento de Pessoal de Nível Superior – Brasil (CAPES), Finance Code 001. Special thanks are extended to Silvia Graziela Torres Miranda for her invaluable technical support, and to Dr. Leandro Licursi for granting access to his autoclave, which was crucial for sterilizing the materials used in our experiments. We are particularly grateful to Dr. Wolfgang Miller for the training provided to Karla Yotoko during her postdoctoral fellowship, funded by CAPES under the program ESENIOR2016289300, focusing on methodologies related to Wolbachia infection studies in drosophilids. Additionally, our gratitude goes to Cristiane Bouzada for her assistance with the English editing of our manuscript, and to Lucas Rosado for creating the Graphical Abstract.

