## Supplementary Material for "Flipping the Script: *Wolbachia* Favoring Males in a Neotropical *Drosophila*"

### Supplementary Material 1
#### Culture Media

##### 1. Banana-Barley Culture Medium

Materials Required

- Corn syrup - 9.5 g
- Banana - 13.8 g
- Distilled Water - 100 ml
- Beer Yeast - 2.8 g
- Barley Flour - 3.0 g
- Agar-agar - 0.8 g
- Methylparaben 0.22 g
- 100 % Ethanol

Procedures

Prepare this medium the day before you plan to use it. Keep it at room temperature. If it´s too wet, flies may sick to it and die.

1. In an Erlenmeyer, place corn syrup, banana, and 50 ml of water.
2. Cook the mixture in a microwave until getting a homogeneous mixture.
3. Add beer yeast, barley flour, agar-agar and the remaining water.
4. Cook long enough to homogenize the ingredients and dissolve the agar.
5. Cool the medium until it is comfortable to the touch.
6. In a separate container, dissolve methylparaben in a small amount of ethanol, then add this to your mixture.
7. Pour the medium into sterilized containers Let it solidify at room temperature.
8. Before using, sprinkle granulated biological yeast on top of the medium.

#### 2. Grape-Fruit Medium

Materials Required

- Agar - 0.8 g
- Grape Juice - 30 ml
- Methylparaben - 0.22 g
- Distilled water - 100 ml
- 100 % Ethanol

Procedures

Prepare this medium a day before use and keep it at room temperature. If it´s too wet, flies may sick to it and die.

1. In a flask, mix agar with 70 milliliters of distilled water.
2. Heat the mixture in a microwave long enough to dissolve the agar.
3. Add the grape juice and heat it again.
4. Cool the medium until it is comfortable to the touch.
5. In a separate container, dissolve methylparaben in a small amount of ethanol, then add this to your mixture.
6. Pour the medium into sterile plates and let it solidify.
7. Before use, add a thin layer of granulated biological yeast, dissolved in water, on top to encourage females' oviposition (egg laying).

### Supplementary Material 2
#### DNA Extraction Protocols

##### Materials Required (for Single-fly or 5-flies Extractions)

- Wizard Genomic DNA Purification Kit - PROMEGA;
- 1.5 ml microcentrifugue tubes;
- Sterilized pestles;
- 0.5 M EDTA (pH 8.0);
- Proteinase K (20 mg/ml);
- water bath (37 °C);
- Isopropanol;
- 70 % Ethanol.
-

##### Single-Fly specifities:

Prepare Tissues

1. Prepare the EDTA/Nuclei Lysis Solution (**20 µl** EDTA / **85 µl** Cell Lysis Solution per sample).
2. Add **100 µl** of the chilled EDTA/Nuclei Lysis Solution. Macerate the sample.
3. Add **2 ul** of Proteinase K and incubate at 37 °C for 30 min.
4. Cool to room temperature.

Lysis and Protein Precipitation

5. Add **34 ul** of Protein Precipitation Solution to the lysate. Vortex and chill on ice for 5 minutes.
6. Centrifugue at 13,000 rpm for 5 minutes.

##### 5-flies specifities:

Prepare Tissues

1. Prepare the EDTA/Nuclei Lysis Solution **(35 µl** EDTA/**150 µl** Cell Lysis solution per sample).
2. Add **180 µl** of the chilled EDTA/Nuclei Lysis Solution. Macerate the sample.
3. Add **5 ul** of Proteinase K and incubate at 37 °C for 30 min.
4. Cool to room temperature.

Lysis and Protein Precipitation

5. Add **60 ul** of Protein Precipitation Solution to the lysate. Vortex and chill on ice for 5 minutes.
6. Centrifugue at 13,000 rpm for 5 minutes.

**DNA Precipitation and Rehydration**

7. Transfer supernatant to a fresh tube containing 600 µl of room temperature isopropanol.
8. Mix gently by turning the tube over for at least 2 minutes.
9. Centrifuge at 13,000 rpm for 5 minutes.
10. Remove supernatant and add 600 µl of room temperature 70 % ethanol.
11. Repeat steps 9 and 10.
12. Air-dry the pellet.
13. Rehydrate the DNA in 20 µl of DNA Rehydration Solution for 1 h at 65 °C or overnight at 4 °C.
14. Add the RNase Solution (**2 µl** for single fly or **5 µl** for 5-flies). Incubate for 1 h at 37 °C.
15. Freeze the samples.

#### Supplementary Material 3
##### Survival Courves

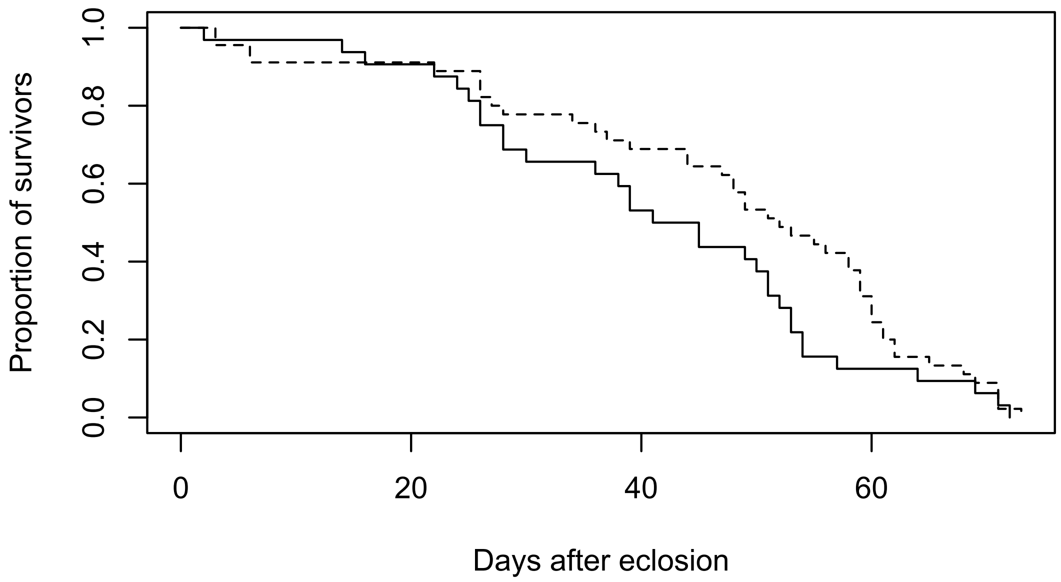


Figure S3.1. Survival of females *Drosophila sturtevanti* (st8 lineage) infected (w+ - solid line) and treated with antibiotic (Rif - dashed line). The Kaplan-Meier survivorship analysis using a log-rank test did not detect a significant difference between the two lineages (p=0.18).

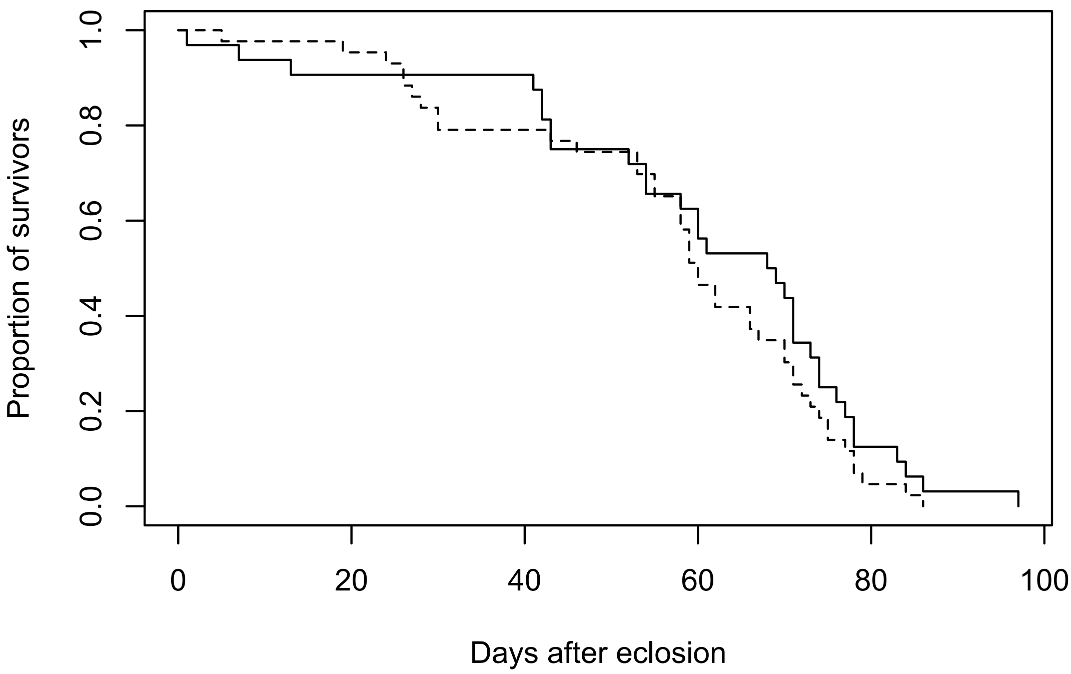


Figure S3.2. Survival of males of *Drosophila sturtevanti* (st8 lineage) infected (w+ - solid line) and treated with antibiotic (Rif - dashed line). The Kaplan-Meier survivorship analysis using a log-rank test did not detect a significant difference between the two lineages (p=0.39).

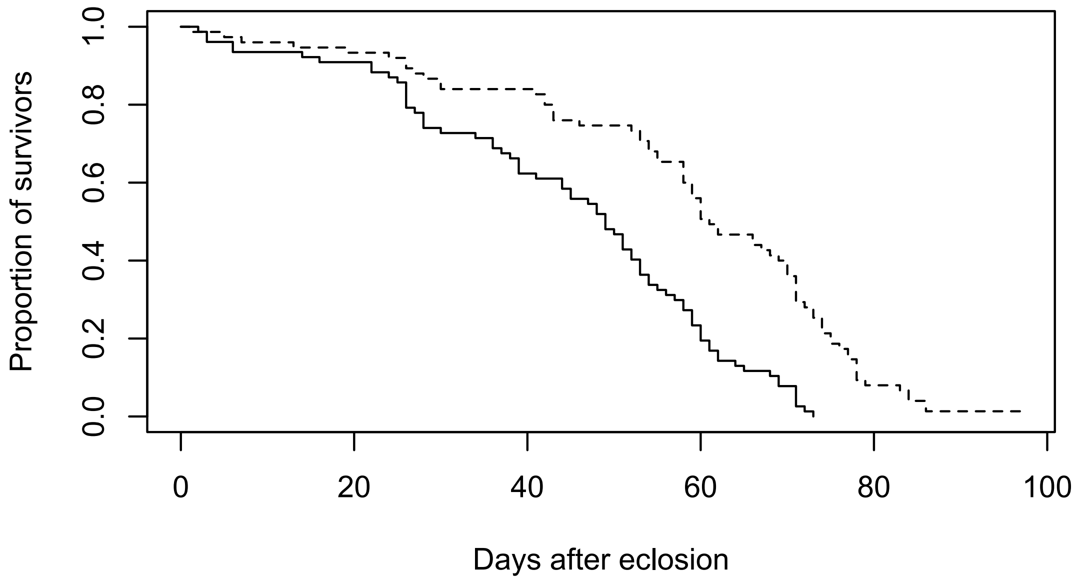


Figure S3.3. Survival of *Drosophila sturtevanti* (lineage st8) females (solid line) and males (dashed line), independent of infected status (infected, w+; or antibiotics-treated, Rif). The Kaplan-Meier survivorship analysis using a log-rank test identified a significant difference between the two sexes (p=0.00011).

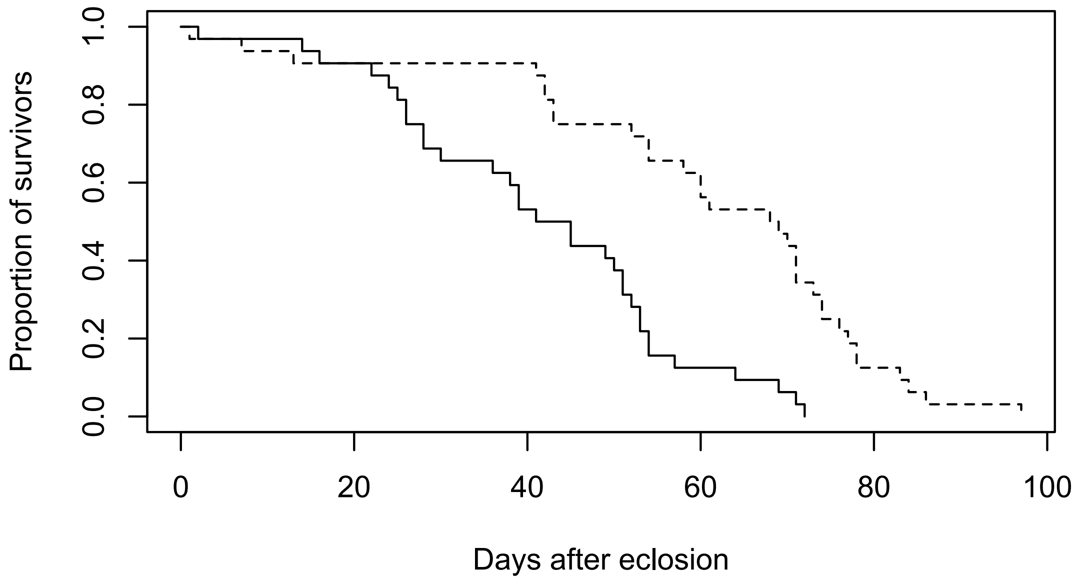


Figure S3.4. Survival of *Wolbachia*-infected (w+) *Drosophila sturtevanti* (lineage st8) females (solid line) and males (dashed line). The Kaplan-Meier survivorship analysis using a log-rank test identified a significant difference between the two sexes (p=0.00026).

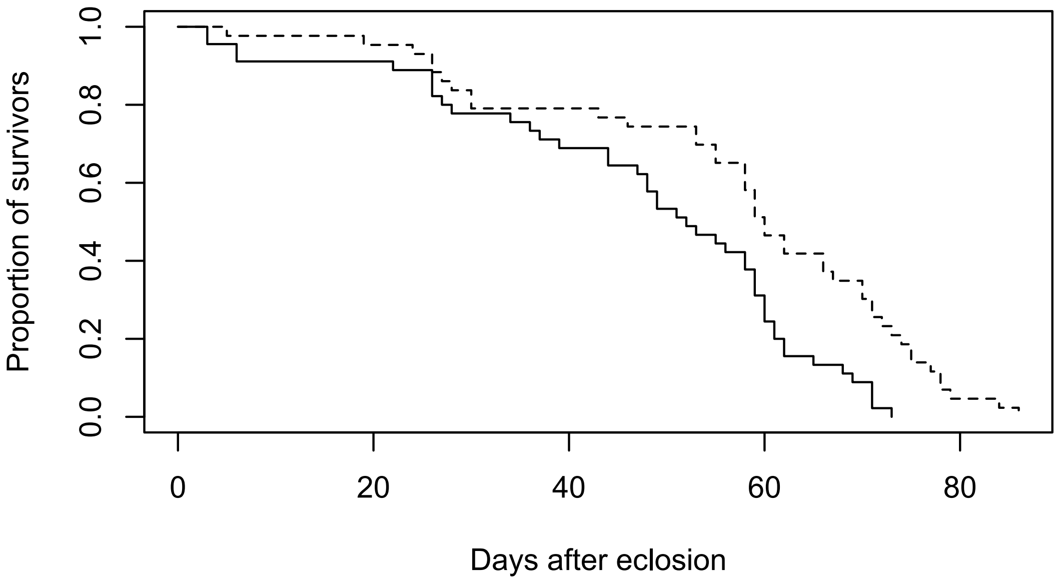


Figure S3.5. Survival of *Drosophila sturtevanti* (lineage st8) Rifampicin-treated (Rif) females (solid line) and males (dashed line). The Kaplan-Meier survivorship analysis using a log-rank test identified a significant difference between the two sexes (p=0.028).
